# Reliability of M1-P15 as a cortical marker for transcallosal inhibition: a preregistered TMS-EEG study

**DOI:** 10.1101/2022.03.10.483631

**Authors:** Agnese Zazio, Guido Barchiesi, Clarissa Ferrari, Eleonora Marcantoni, Marta Bortoletto

**Affiliations:** Neurophysiology Lab, IRCCS Istituto Centro San Giovanni di Dio Fatebenefratelli, Brescia (Italy); Cognition in Action (CIA) Unit, PHILAB, Milan (Italy); Department of Philosophy, Università degli Studi di Milano, Milan (Italy); Statistics Unit, IRCCS Istituto Centro San Giovanni di Dio Fatebenefratelli, Brescia (Italy)

**Keywords:** TEPs, effective connectivity, interhemispheric inhibition, motor system, ipsilateral silent period, bimanual coordination

## Abstract

**Background:** In a recently published study combining transcranial magnetic stimulation and electroencephalography (TMS-EEG), we provided first evidence of M1-P15, an early component of TMS-evoked potentials, as a measure of transcallosal inhibition between motor cortices. However, considering the technical challenges of TMS-EEG recordings, further evidence is needed before M1-P15 can be considered a reliable index.

**Objective:** Here, we aimed at validating M1-P15 as a cortical index of transcallosal inhibition, by replicating previous findings on its relationship with the ipsilateral silent period (iSP) and with performance in bimanual coordination. Moreover, we aimed at inducing a task-dependent modulation of transcallosal inhibition.

**Methods:** A new sample of 32 healthy right-handed participants underwent behavioral motor tasks and TMS-EEG recording, in which left and right M1 were stimulated during bimanual tasks and during an iSP paradigm. Hypotheses and methods were preregistered before data collection.

**Results:** We successfully replicated our previous findings on the positive relationship between M1-P15 amplitude and the iSP normalized area. However, we did not confirm the relationship between M1-P15 latency and bimanual coordination. Finally, we show a task-dependent modulation of M1-P15 amplitude, which was affected by the characteristics of the bimanual task the participants were performing, but not by the contralateral hand activity during the iSP paradigm.

**Conclusions:** The present results corroborate our previous findings in validating the M1-P15 as a reliable cortical marker of transcallosal inhibition, and provide novel evidence of its task-dependent modulation. Importantly, we demonstrate the feasibility of a preregistration approach in the TMS-EEG field to increase methodological rigor and transparency.

## Introduction

The combination of transcranial magnetic stimulation and electroencephalography (TMS-EEG) provides a unique perspective on effective connectivity [1,2], defined as the description of causal relationships between brain areas [3]. Pioneering studies have shown that TMS-evoked potentials (TEPs) represent the propagation of cortical responses from the stimulated area to the connected ones [4,5], conveying state-dependent information [6,7]. In recent years, a deeper understanding of the TMS-EEG signal was ensured by the integration with other neuroimaging techniques, such as the magnetic resonance, indicating that TMS signal propagates mainly on structural connections of the stimulated networks [8–10].

Several studies have employed TEPs to measure interhemispheric communication, and specifically transcallosal connections between motor cortices. A first measure of interhemispheric signal propagation has been obtained as the ratio between TEPs recorded on EEG channels over the hemisphere contralateral to TMS and on the ones over the stimulated hemisphere, averaging over a time window of about 100 ms after the TMS pulse. This approach has been firstly described by Voineskos and colleagues [11], who showed an inverse relationship between this measure and the microstructural integrity of the corpus callosum. Subsequent studies on healthy participants have further explored this index to unravel its relationship with a peripheral measure derived from the double-coil paradigm [12] as well as the underlying neurochemical mechanisms [13]. Furthermore, it has been also applied to study the interhemispheric signal propagation in clinical populations, such as patients with stroke [14] and autism spectrum disorder [15].

Despite this approach first described by Voineskos and coworkers [11] has been proven to be informative as a global measure of interhemispheric signal propagation, it is known that the transcallosal conduction delay (TCD), i.e. the timing of interhemispheric connectivity along the fibers of the corpus callosum, is much faster and occurs within 20 ms in the motor system [16,17]. In this context, preliminary TMS-EEG evidence had suggested that TEPs can provide information on TCD, showing that after primary motor cortex (M1) stimulation the signal is transferred to the contralateral motor areas in the first tens of ms [4,18]. Therefore, we have recently proposed to study TCD by measuring early TEPs occurring within 20 ms after theTMS pulse, exploiting the excellent temporal resolution of EEG.

In our recent study [19], we combined TMS-EEG with diffusion tensor imaging (DTI), which provides microstructural information on callosal integrity, and with the ipsilateral silent period (iSP), a well-known peripheral measure of interhemispheric inhibition obtained from TMS and electromyography [20]. Our results provided first evidence of M1-P15, a positive component occurring approximately 15 ms after M1 stimulation, as a TEP-derived measure of transcallosal inhibition between motor cortices. Indeed, the latency of M1-P15 was predicted by DTI structural connectivity, such that the higher the diffusivity along the fibers of the body of the corpus callosum, the shorter the latency of M1-P15. This first evidence suggested that M1-P15 latency may be considered a measure of TCD. Moreover, our findings indicated that the amplitude of M1-P15 reflects the strength of the transcallosal inhibition, as shown by a positive relationship with the magnitude of the iSP. Finally, the TCD as indexed by M1-P15 latency was associated with bimanual coordination performance, such that shorter left-to-right together with longer right-to-left TCD was associated with better temporal performance in bimanual finger opposition movements.

Considering the technical challenges inherent in the study of early EEG responses to TMS, further evidence is needed to make the M1-P15 a valid and reliable measure. Moreover, the relationship between M1-P15 and bimanual coordination required further investigation. First, despite a relationship between M1-P15 latency and behavioral performance was expected based on previous theories [21], the interaction with the direction of information flow between hemispheres was not. Second, in the original study the M1-P15 and the bimanual performance were recorded separately and under different conditions, leaving an open question on whether the M1-P15 could be recorded also during the bimanual task and whether its latency was predictive of bimanual performance. Finally, another unexplored topic regards the possibility of modulating transcallosal inhibition as indexed by M1-P15.

Based on previous findings, we designed a new TMS-EEG study with the following main aims and hypotheses:

i. Validating M1-P15 as a measure of transcallosal inhibition: we expect to replicate the positive relationship between M1-P15 amplitude and the magnitude of iSP;
ii. Evaluating the behavioral relevance of M1-P15: we expect to replicate the relationships between M1-P15 latency and bimanual coordination during the sequential thumb-to-finger opposition task; if the replication was successful, we aimed at exploring the potential cost of TCD asymmetry on unimanual performance;
iii. Inducing a modulation of the transcallosal inhibition, by manipulating the activity of the hand contralateral to the stimulation.

Our hypotheses, as well as the methods and the analyses we planned to run, were preregistered on OSF before data collection (https://osf.io/pg78j/), with the aim of increasing the transparency of the research process and of providing an unbiased picture of the results, thus contributing in improving scientific replicability [22].

## Materials and methods

### Participants

Thirty-two right-handed healthy participants were enrolled in the study after giving written informed consent (19 women; mean age ± SD: 29.7 ± 8.6 years, range: 19-48 years). Participants who took part in the study by Bortoletto et al. (2021) were not enrolled in the present study. They had no history of neurological disorders nor contraindications to TMS [23]. The study was performed in accordance with the ethical standards of the Declaration of Helsinki and approved by the Ethical Committee of the IRCCS Istituto Centro San Giovanni di Dio Fatebenefratelli (Brescia), where the experiment took place. One participant did not complete all the experimental blocks, leaving 31 subjects for the off-line analyses. In the off-line analyses, two additional participants were excluded due to the presence of residual artifacts, one in TMS-EEG and one in the iSP preprocessing, respectively.

### Design and procedure

Participants have been involved in a within-subject single-session design experiment (Figure 1). They were comfortably seated in front of a computer monitor with their forearms leaning on a desk. The experiment involved the following steps: behavioral motor tasks, TMS-EEG recording during bimanual tasks and TMS-EEG recording during an iSP paradigm. The total duration of the experiment was about 2.5 hours.

**Figure 1.**
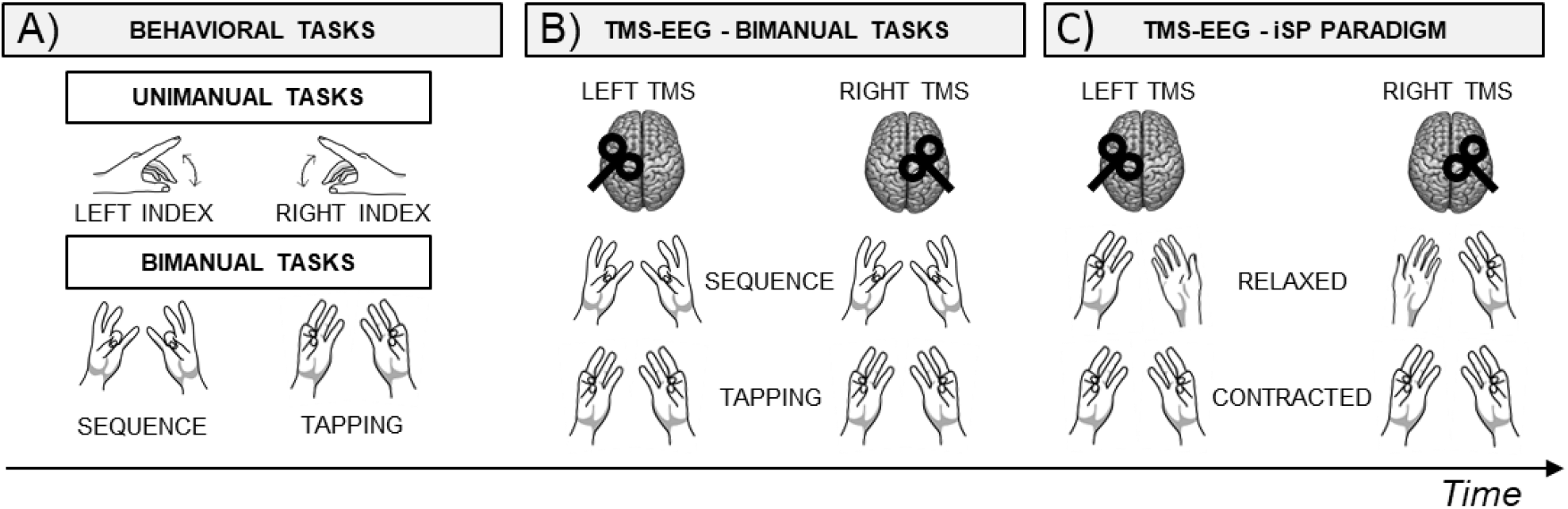
Experimental design. Schematic representation of the within-subject single-session design experiment. Block order was counterbalanced among participants using a latin square design. **A)** First, participants performed the behavioral tasks, including a unimanual task performed with the left and the right hand, followed by two bimanual tasks, i.e., *Sequence* and *Tapping*. **B)** Then, participants performed the *Sequence* and the *Tapping* tasks during TMS-EEG recording, after stimulation of left and right M1, in separate blocks. **C)** Finally, participants performed an iSP paradigm during TMS-EEG recording after stimulation of left and right M1, in separate blocks. Specifically, in the iSP paradigm the APB muscle of the hand ipsilateral to TMS was always contracted, while the contralateral APB was *Relaxed* or *Contracted*, in separate blocks.

After EEG and EMG montage, the motor hotspot for APB muscle was localized as the scalp site eliciting the highest and most reliable motor-evoked potentials (MEPs) with the same TMS intensity. Coil orientation was kept approximately 45° from the midline, inducing an anterior-to-posterior and posterior-to-anterior (AP-PA) current direction in the brain. Then, the resting motor threshold (rMT) was estimated using the maximum-likelihood threshold hunting algorithm [24,25], a variant of the best parameter estimation by sequential testing (best PEST) procedure [26]. This procedure was performed for each hemisphere.

#### Behavioral tasks

Behavioral tasks consisted of an unimanual task (see Supplementary Materials) and two bimanual coordination tasks (Figure 1A). Participants wore disposable gloves; then, to measure bimanual performance, a conductive sensor was applied on each of the participants’ finger tips with a double sided tape. In this way, sensors could be adjusted according to hand size and shape.

The bimanual coordination tasks involved a metronome-paced in-phase movements at 2 Hz: a repetitive mirror-symmetrical thumb-to-index opposition task (*Tapping*) and a sequential mirror-symmetrical thumb-to-finger opposition task (*Sequence*), as in [19,27]. Specifically, in the *Sequence* task participants were asked to oppose their thumb to the other fingers, in the following order: index, middle, ring and little finger. Participants performed 3 blocks for each bimanual task; each block lasted about 1 minute. The metronome sound was presented through the earphones, overlapping with a white noise. The white noise was added to make the bimanual tasks comparable in the behavioral tasks and in the TMS-EEG recording.

#### TMS-EEG recording

Single biphasic TMS pulses were delivered over left (LTMS) and right (RTMS) M1 using a Magstim Rapid^2^ stimulator connected to an Alpha B.I. Coil Range 70 mm (Magstim Company, Whitland, UK), while EEG was continuously recorded with a TMS-compatible system (g.HIamp, g.tec medical engineering GmbH, Schiedlberg, Austria). TMS intensity was set at 110% of the average rMT estimated for the left and the right hemisphere, as in our previous study [19] (mean rMT ± SE: LTMS, 62.6 ± 2.1; RTMS: 62.8 ± 2.0). The charge delay was set at 350 ms and coil position was monitored with a neuronavigation system (Softaxic 3.4.0; EMS, Bologna, Italy). To attenuate the contamination of TEPs with sensory artifacts, a thin layer of foam was applied on the coil, and participants wore noise-canceling earphones playing white noise (the volume was individually adjusted to mask the TMS click but avoiding earing discomfort). EEG was recorded by using 74 passive Ag/AgCl electrodes with 10/10 international system; the reference was placed on FPz and the ground on the nose (EasyCap, BrainProducts GmbH, Munich, Germany). Skin/electrode impedance was kept below 5 kΩ. EMG was recorded with a bipolar belly-tendon montage on left and right APB (Ag/AgCl pre-gelled surface electrodes, Friendship Medical, Xi’an, China). Sampling rate was set at 9600 Hz and no filters were applied in recording.

During the bimanual tasks (Figure 1B), the experimental blocks were the same as in the behavioral tasks (i.e., 3 blocks for *Tapping* and 3 blocks for *Sequence*). TMS pulses were randomly delivered at 30 different time intervals (between 16 and 500 ms; see Pilot Study in Supplementary Materials) after the metronome sound, with an inter-stimulus interval between TMS pulses of at least 1 s. The order of the stimulated hemispheres as well as of the tasks was counterbalanced among participants with a Latin square design.

At the end of the bimanual tasks, sensors were removed from the participants’ fingers. Instead, a pressure sensor was applied on the second phalanx of their left and right index finger. Participants were asked to press on the sensor with their thumb, thus inducing a contraction of APB muscle (Figure 1C). Specifically, in the *Contracted* conditions, they were asked to press on the sensors with both hands, while in the *Relaxed* conditions they had to press on the sensor only with their hand ipsilateral to TMS, keeping the contralateral hand relaxed. Participants were asked to press between 40% and 60% of their maximal strength (previously assessed for each hand within a time interval of 10 s). Every time the pressure level fell out of the required range, a visual feedback appeared on the screen indicating the actual pressure compared to the target range. Once the pressure level was reached, visual feedback disappeared and only the fixation cross remained in the center of the screen, to avoid eye movements during TMS-EEG recording. TMS was delivered only when the pressure on the sensors was within the target range.

### Analysis

#### Behavioral performance

Performance in the bimanual tasks was evaluated as the interhand interval (i.e., unsigned time difference in ms) between the onset of finger tap with the left hand and onset of the corresponding finger tap with the right hand (interhand interval values longer than 150 ms were excluded, as well as the ones exceeding 2 SD within each subject [19]). Since we calculated the absolute value of interhand interval, data were log-transformed to avoid skewness.

#### TEPs

TEPs data analysis was performed in MATLAB R2020b (The Mathworks, Natick, MA, USA) with custom scripts using EEGLAB v.2020.0 [28] and FieldTrip [29] functions. If not otherwise specified, default parameters for EEGLAB and FieldTrip function were used. For each participant, the first pre-processing steps merged in the same dataset the two conditions of interest with the same stimulation site: for example, the conditions LTMS-*Contracted* and *LTMS-Relaxed* were analyzed as one dataset (the same for RTMS-*Contracted* and RTMS-*Relaxed*, LTMS-*Tapping* and LTMS-*Sequence*, RTMS-*Tapping* and RTMS-*Sequence*). This approach minimized the risk that differences between conditions that we compared could arise from dissimilarities in the preprocessing steps. Continuous TMS-EEG data was interpolated for 3 ms around the TMS trigger, high-pass filtered at 1 Hz (windowed sinc FIR filter using EEGLAB function ‘pop_eegfiltnew’, order 31681), downsampled at 4800 Hz and epoched from 200 ms before to 500 ms after the TMS pulse. Then, the source-estimate-utilizing noise-discarding (SOUND) algorithm [30]; spherical 3-layer model, regularization parameter: λ=.01) was applied to discard noise measurement, and a first artifact rejection was performed to discard highly artifactual trials based on visual inspection. After SOUND, we run an independent component analysis (ICA) for ocular artifact correction using the EEGLAB function ‘pop_runica’ (infomax algorithm, 73 channels included, 73 ICA components computed; components relative to horizontal and vertical ocular movements are visually inspected and discarded). Then, we applied the signal-space projection and source-informed reconstruction (SSP-SIR) algorithm [31] for TMS-evoked muscular artifact removal (correction in the first 50 ms after TMS pulse). Principal components were visually inspected and discarded if they represented a high-frequency signal time-locked the TMS pulse; this step has been performed by two independent researchers. Finally, after a low-pass filter at 70 Hz (IIR Butterworth filter, order 4, using the EEGLAB function ‘pop_basicfilter’), data was visually inspected as a final check to discard trials with residual artifacts, re-reference to the linked mastoids, epoched from −100 to 400 ms, baseline corrected in the 100 ms preceding the TMS, and averaged according to the experimental condition (Figure 2). On average, 97.5% trials per condition were considered. The analysis steps followed the steps of the preregistered pipeline (https://osf.io/pg78j/), except for the high-pass filter on the continuous TMS-EEG data, which was modified into 1 Hz instead of 0.1 Hz for computational demands. Although in the present work we were not interested in slow potentials, we preliminary checked on the Pilot data that high-pass filtered TEPs at 1 Hz or 0.1 Hz were qualitatively comparable (Figure S2).

**Figure 2.**
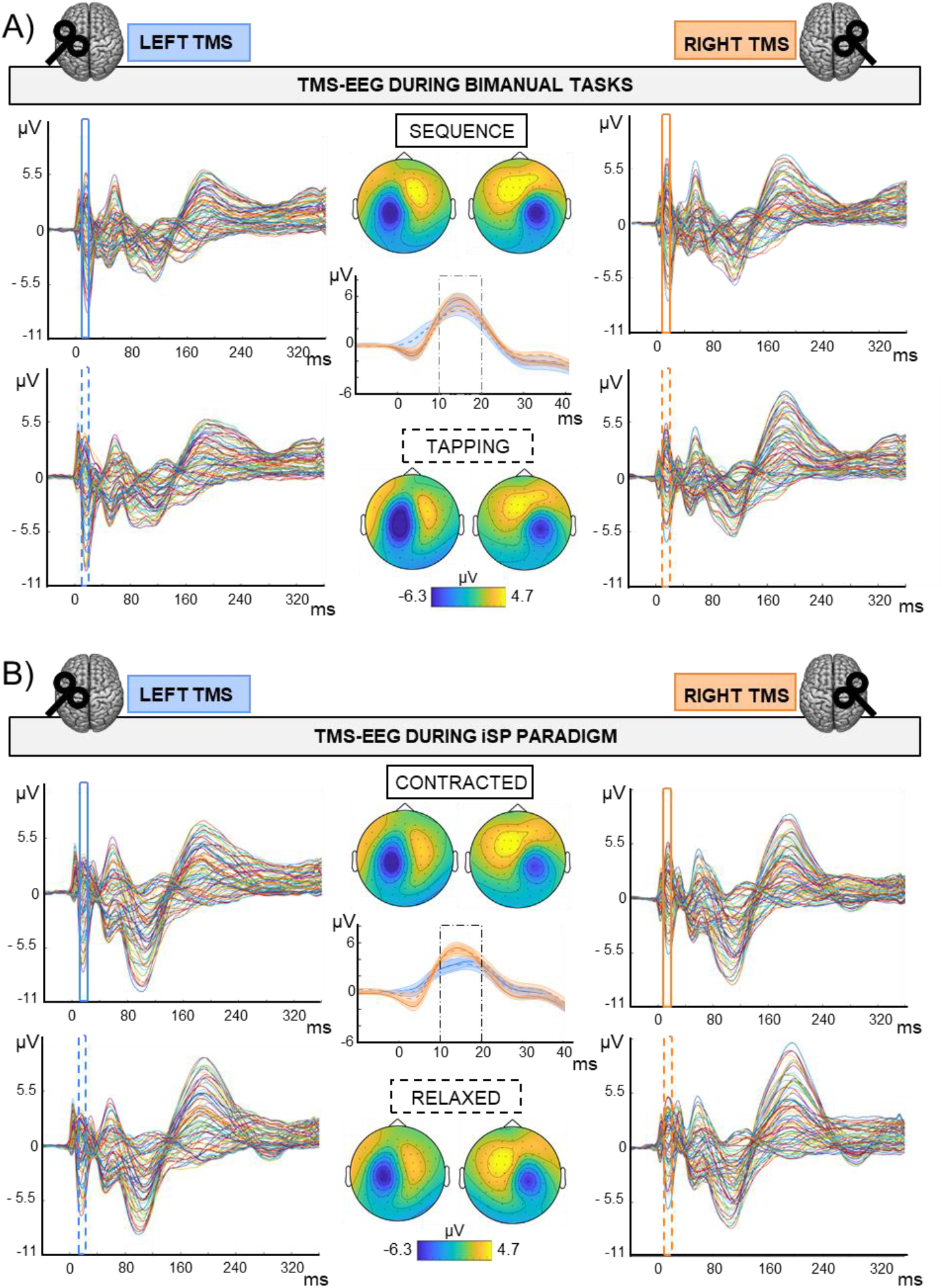
TMS-EEG output in the different experimental conditions. Grand average (N=30) of TEPs and M1-P15 topographies obtained during the bimanual tasks (A) and the iSP paradigm (B). M1-P15 topographies were obtained by averaging over time between 10 and 20 ms after the TMS pulse; amplitude range as indicated in colorbars. Central panels show M1-P15 on the average of F4-FC4 channels after LTMS (blue), and F3-FC3 after RTMS (orange); SE in shaded error bars. **A)** 2 × 2 Hemisphere (LTMS, RTMS) X Task (*Sequence, Tapping*) design; Top row and continuous traces: TMS-EEG data during *Sequence*; bottom row and dashed traces: TMS-EEG data during *Tapping*. **B)** 2 × 2 Hemisphere (LTMS, RTMS) X Contraction (*Contracted, Relaxed*) design; Top row and continuous traces: TMS-EEG data during *Contracted*; bottom row and dashed traces: TMS-EEG data during *Relaxed*.

The M1-P15 identification was performed as follows. For each subject we first averaged over all conditions following the stimulation of the same hemisphere to increase the signal-to-noise ratio (i.e., *Tapping, Sequence, Contracted* and *Relaxed* during LTMS, and *Tapping, Sequence, Contracted* and *Relaxed* during RTMS). From the grand-average of all participants we identified the channels contralateral to TMS with the highest amplitude in the time window between 7 and 25 ms: F4-FC4 for LTMS, F3-FC3 for RTMS. Then, we identified the maximum on the pooled signal of the two channels in the time window between 7 and 25 ms; in case more than one peak was present or no peak could be detected, we manually restricted the time-window to match the M1-P15 topography (i.e., contralateral positivity over frontocentral electrodes). Then, we divided the data into the different experimental conditions, and the M1-P15 peak was automatically detected within 10 ms around the individual peak identified previously. M1-P15 amplitude was calculated by averaging over 10 ms around the peak. Overall, this procedure allowed us to perform a semi-automatic peak detection, avoiding subjective bias between the experimental conditions that we aim to compare.

#### iSP

The first preprocessing steps for the EMG trace ipsilateral to TMS were the same as for TMS-EEG data (i.e., interpolation, high-pass filter, downsample and epoching). Then, EMG was band-pass filtered between 100 and 1000 Hz (default FieldTrip filters) and rectified. For each participant, for each trial we checked whether the signal was lower than the mean of prestimulus baseline (calculated between 150 and 50 ms before the TMS pulse) for at least 10 ms in the time window between 25 and 75 ms after the TMS pulse; trials which did not satisfy this criterion were discarded. On average, 58.5 trials (64.9%) were considered in the average of each condition (range: 26-87 trials). Then, for each participant and condition we averaged over trials, and calculated the iSP with the same parameters as the ones applied on single trials. One subject was excluded in this step because no iSP was detected, leaving 29 participants for the statistical analyses involving the iSP. Finally, we calculated the normalized iSP area using the following formula [(area of the rectangle defined as mean EMG x iSP duration)-(area underneath the iSP)]. Then, the normalized iSP area was defined as the ratio between the iSP area and the area underneath EMG from −150 to 50 ms preceding TMS [19,32]. Even if the number of trials considered for iSP calculation are consistent with the iSP literature, they represent only a subset of the trials presented and that were considered for M1-P15 calculation. To check whether the relationship between M1-P15 amplitude and iSP normalized area was maintained even when considering a comparable number of trials for the iSP estimation, in a control analysis we calculated the iSP by using 5 ms as minimum duration, both on single trials and on the average, leaving on average 86.4 trials for each condition (96%, range 75-90).

### Statistical analysis

i. To validate M1-P15 as a measure of interhemispheric inhibition, we run a linear mixed model (LMM) with random slopes and intercepts [33], testing whether M1-P15 amplitude (repeated independent variable) predicted the iSP normalized area (repeated dependent variable), as defined in the preregistration. The LMM had Condition (4 levels: *LTMS-Contracted, LTMS-Relaxed, RTMS-Contracted, RTMS-Relaxed*) as fixed factor and Subject as random factor, with each condition repeated within subjects. We also performed two explorative (i.e., not defined in the preregistration) control analyses to exclude that the relationship between M1-P15 amplitude and the iSP normalized area could be explained by a common factor, namely the TMS intensity. Specifically, we assessed the relationship between rMT averaged between hemispheres (as predictor) and the M1-P15 amplitude and iSP normalized area (as dependent variables, in separate models). We run LMMs with random slopes and intercepts with Condition (4 levels: *LTMS-Contracted, LTMS-Relaxed, RTMS-Contracted, RTMS-Relaxed*) as fixed factor and Subject as random factor, with each condition repeated within subjects.
ii. LMMs with random slopes and intercepts [33] were also applied to assess the behavioral relevance of M1-P15, as planned in the preregistration. Bimanual coordination performance in the *Sequence* task was set as dependent variable, using the interhand interval of each Tap of the bimanual task as fixed effect and Subject as a random effect (with taps repeated within subjects). For each subject, we considered an equal number of taps, by selecting the first N taps (where N is the number of taps of the participant with the minimum number of taps; N= 268). As a predictor, we first considered the M1-P15 latency measured during the iSP paradigm. We run separate LMMs considering three measures of M1-P15 latency as predictor, namely: the mean value of *Contracted* and *Relaxed* conditions in the LTMS blocks, the mean value of *Contracted* and *Relaxed* conditions in the RTMS blocks and in the ratio between the two. Then, we performed the same statistical models but considering as predictor the M1-P15 latency recorded while participants were performing the bimanual *Sequence* task. Since no significant relationships were observed between M1-P15 latency and bimanual performance in the *Sequence* task (see Results), we did not performed the preregistered analyses on the relationships between M1-P15 and bimanual performance in the *Tapping* task, as well as between M1-P15 and unimanual performance. Besides the preregistered tests, we also ran additional control analyses on the behavioral tasks. We compared the bimanual performance between *Sequence* and *Tapping* by means of a two-tailed t-test for dependent samples, and we checked for an effect of time by comparing the bimanual performance in the three blocks, as indexed by the interhand interval measured during *Sequence*, by means of one-way repeated-measures analysis of variance (rm-ANOVA) with the 3-level factor Block. We also explored possible modulations of M1-P15 latency, both during the bimanual tasks (2 × 2 rm-ANOVAs with factors Hemisphere and Task) and during the iSP paradigm (2 × 2 rm-ANOVAs with factors Hemisphere and Contraction).
iii. To assess the modulation of the interhemispheric inhibition, we first tested the effects of contraction levels of the hand contralateral to the stimulation. As defined in the preregistered analyses, we performed a semi-parametric repeated-measures multivariate analysis of variance (rm-MANOVA; 10000 iterations, modified ANOVA-type statistic – MATS, parametric bootstrap approach for resampling) on M1-P15 amplitude and iSP normalized area (as dependent variables), with Contraction (*Contracted, Relaxed*) and Hemisphere (*LTMS, RTMS*) as predictors. Post-hoc analyses were performed by computing separate 2 × 2 rm-ANOVA on M1-P15 amplitude and iSP. Furthermore, we explored whether M1-P15 amplitude was modulated by the bimanual task performed during the TMS-EEG recording, by running a 2 × 2 rm-ANOVA with factors Hemisphere (*LTMS, RTMS*) and Task (*Sequence*, *Tapping*). Finally, we compared the rMT for the left and the right hemisphere by means of a two-tailed dependent sample t-test.

Statistical significance was set at *p* < 0.05. LMMs and rm-MANOVA were run in R software v. 4.0.0 [34] using the lme4 and the MANOVA.RM package, respectively, and remaining statistical analyses were performed using jamovi v. 1.6.15 [35]. If not otherwise specified, mean ± SE is reported in parentheses.

## Results

i. As described below, results successfully replicated our previous findings, thus corroborating the evidence of M1-P15 as a cortical index of interhemispheric inhibition. LMMs showed that M1-P15 amplitude positively predicted the normalized area of the iSP (*t* = 3.3, *p* = 0.001), so that the larger the M1-P15, the stronger the interhemispheric inhibition in the ipsilateral APB (Figure 3A). Similar results were obtained by calculating the iSP with a minimum duration of 5 ms instead of 10 ms (*t* = 2.85, *p* = 0.006). Moreover, LMMs showed a non-significant relationship between rMT and P15 amplitude (*t_(29)_* = 1.26, *p* = 0.22; Figure 3B) and between rMT and iSP normalized area (*t_(28_*) = 0.83, *p* = 0.42; Figure 3C), suggesting that the significant relationship between the M1-P15 amplitude and the magnitude of the iSP cannot be explained by a third common factor, namely the rMT, which determines the TMS intensity used throughout the experiment.
ii. In contrast with our previous work, LMMs did not reveal any significant relationship between M1 - P15 latency recorded during the iSP paradigm and bimanual performance in the *Sequence* task, indexed by the interhand interval. No significant effects were observed for LTMS (*t* = 0.74, *p* = 0.47), RTMS (*t* = 1.9, *p* = 0.07), nor the ratio between the two (*t* = −0.42, *p* = 0.69; Figure 4). Null effects were observed also when considering the M1-P15 latency measured while participants were performing the *Sequence* task (LTMS: *t =* −0.09, *p* = 0.93; RTMS: *t* = 0.74, *p* = 0.48; rate: *t* = −0.76, *p* = 0.46). From the control analysis on the behavioral measure of performance, we excluded an effect of fatigue or learning throughout the behavioral task, because one-way rm-ANOVA with the 3-level factor Block on the interhand interval showed a non-significant effect of time (*F_(2,58)_* = 0.75, *p* = 0.48). Overall, the M1-P15 latency was unaffected by the factors we manipulated, i.e. Hemisphere, both during bimanual paradigm (*F_(1,29)_* = 0.02, *p* = 0.9) and during iSP paradigm (*F_(1,28)_* = 2.13, *p* = 1.56), Task during bimanual paradigm (*F_(1,29)_* = 0.56, *p* = 0.46) and Contraction during iSP paradigm (*F_(1,28)_* = 0.47, *p* = 0.5); no interaction were observed (Hemisphere X Task: *F_(1,29)_* = 1.91, *p* = 0.18; Hemisphere X Contraction: *F_(1,28)_* = 0.01, *p* = 0.92).
iii. The strength of interhemispheric inhibition did not appear to be affected by the contraction levels of contralateral hand activity during the iSP paradigm; however, it was effectively modulated by the bimanual task that participants were performing during the TMS-EEG recording. Regarding the data collected during the iSP paradigm, the rm-MANOVA revealed no significant main effect of Contraction (MATS*_(2)_* = 0.64, *p* = 0.22) nor Contraction X Hemisphere (MATS*_(2)_* = 1.00, *p* = 0.53) on the iSP normalized area and M1-P15 amplitude. We observed a main effect of Hemisphere (MATS*_(2)_* = 6.66, *p* = 0.023); post-hoc analyses by means of rm-ANOVAs revealed a significant main effect of Hemisphere on iSP only (*F_(1,28)_* = 4.24, *p* = 0.049), with a larger iSP after RTMS compared to LTMS; this effect did not reach the level of significance on M1-P15 amplitude (*F_(1,28_*) = 3.69, *p* = 0.065). The control analysis indicated that left and right rMT did not differ (*t_(29)_* = 0.11, *p* = 0.91), suggesting that the effect of Hemisphere cannot be explained by a difference in TMS intensity. Conversely, on the TMS-EEG data collected during the bimanual tasks, the 2 × 2 rmANOVA on M1-P15 amplitude showed a significant main effect of Task (*F_(1,29)_* = 6.43, *p* = 0.017), explained by a larger M1-P15 recorded during *Sequence* (5.17 ± 0.5 μV) compared to *Tapping* (4.44 ± 0.5 μV; Figure 5A). No significant main effect of Hemisphere (*F_(1,29)_* = 0.03, *p* = 0.87) nor interaction Task X Hemisphere (*F_(1,29)_* = 1.79, *p* = 0.19) were observed. Interestingly, the interhand interval was significantly higher (i.e., lower bimanual coordination) in the *Sequence* (25.7 ± 8.6 ms) compared to the *Tapping* task (15.5 ± 5.1 ms; *t_(29)_* = 8.07, *p* < 0.001; Figure 5B).

**Figure 3.**
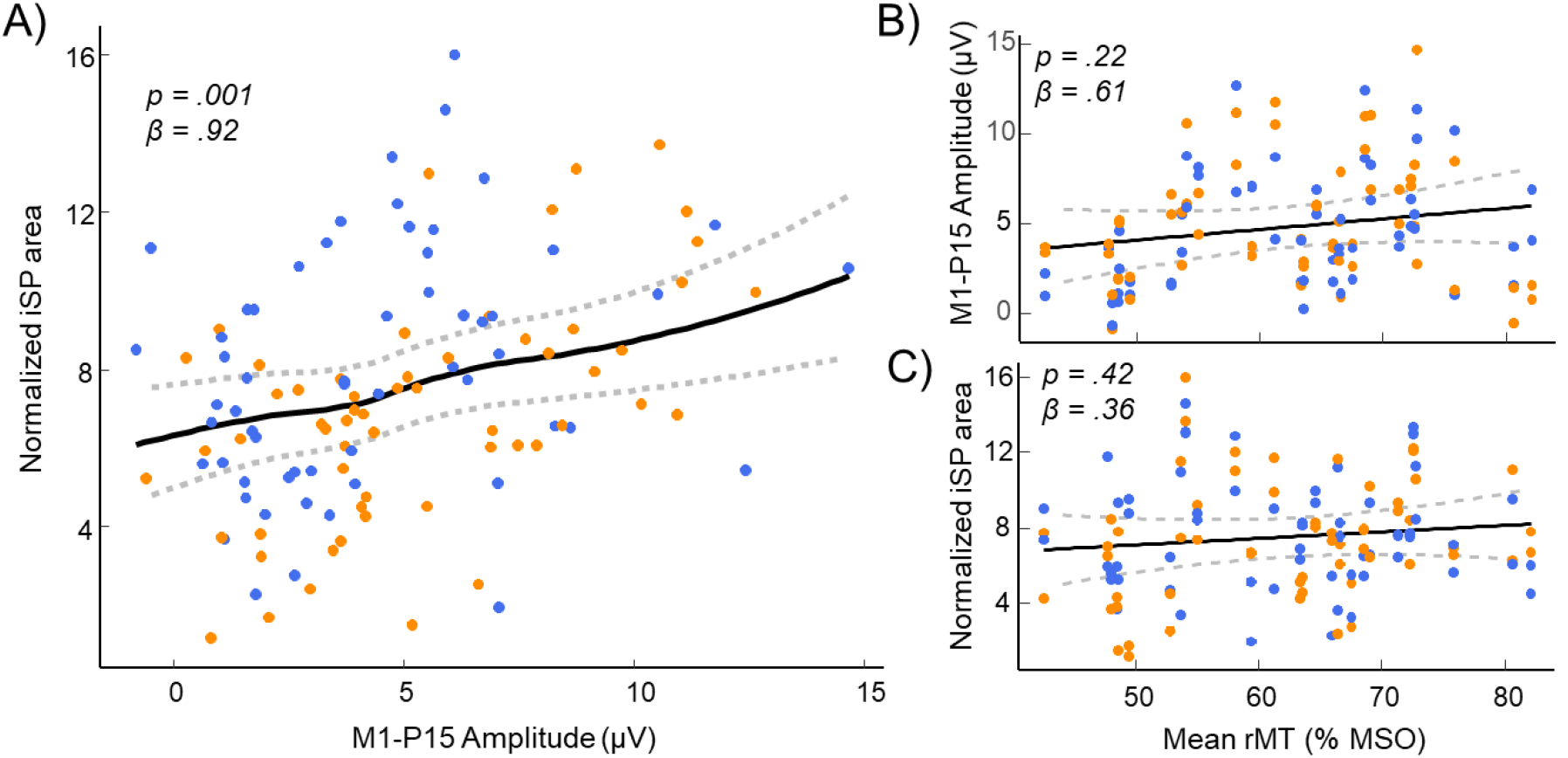
M1-P15 as a measure of transcallosal inhibition. **A)** Significant positive relationship between M1-P15 amplitude and normalized iSP area, indicating that the larger the M1-P15 amplitude, the greater the iSP. **B-C)** Results of the control analyses showing non significant relationships between the mean rMT of both hemispheres and the M1-P15 amplitude (B) and the normalized iSP area (C). Blue dots indicate LTMS, orange dots indicate RTMS. Fitted curves were drawn by applying a smoothed spline to predicted values in the LMMs obtained by a bootstrap procedure (n = 500 simulations); dashed lines represent the 95% confidence interval.

**Figure 4.**
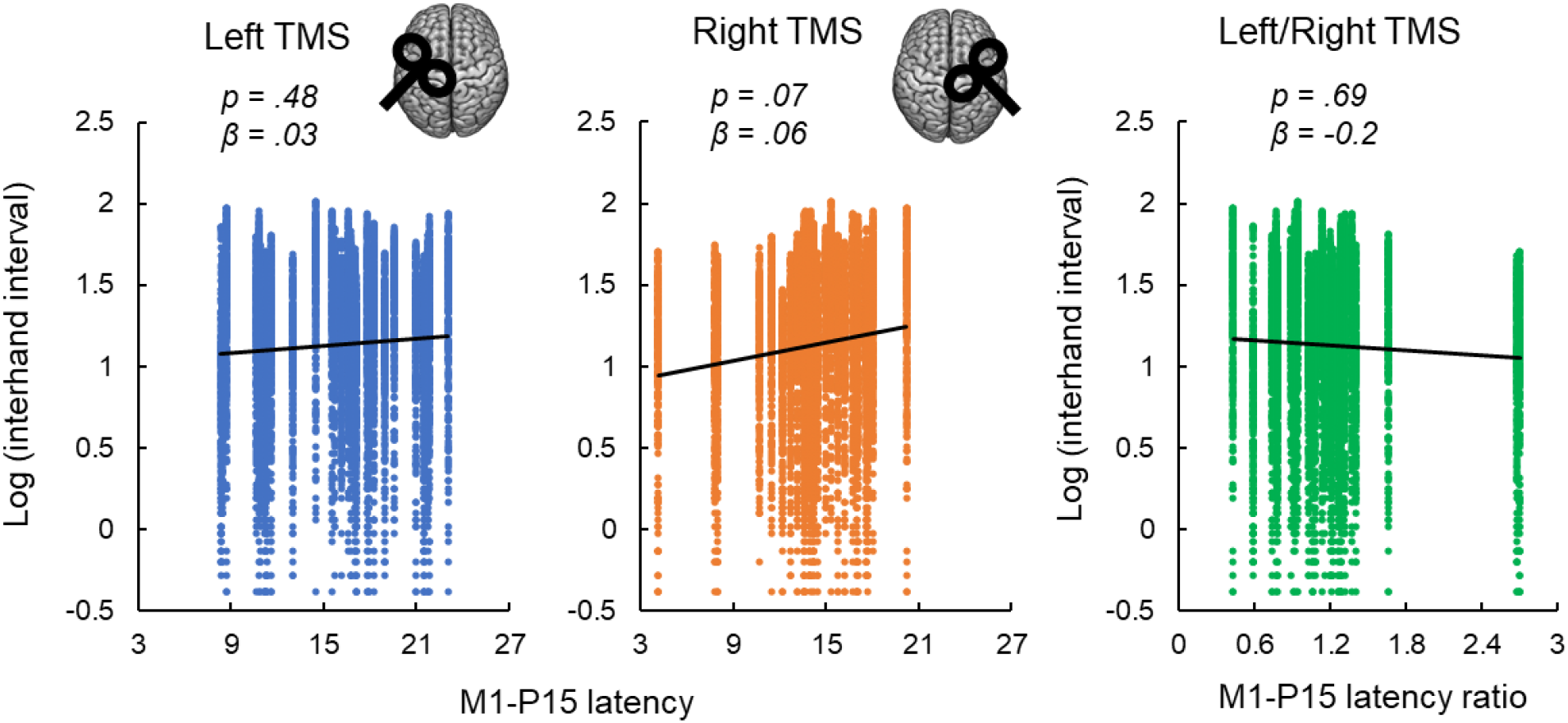
Non-replication of the relationships between M1-P15 latency and bimanual performance. M1-P15 latency recorded during the iSP paradigm was not predictive of bimanual performance in the *Sequence* task, neither after LTMS (left panel), nor after RTMS (central panel) nor the ratio between LTMS and RTMS, as revealed by LMM considering single-trial interhand intervals. Fitted lines represent linear trends.

**Figure 5.**
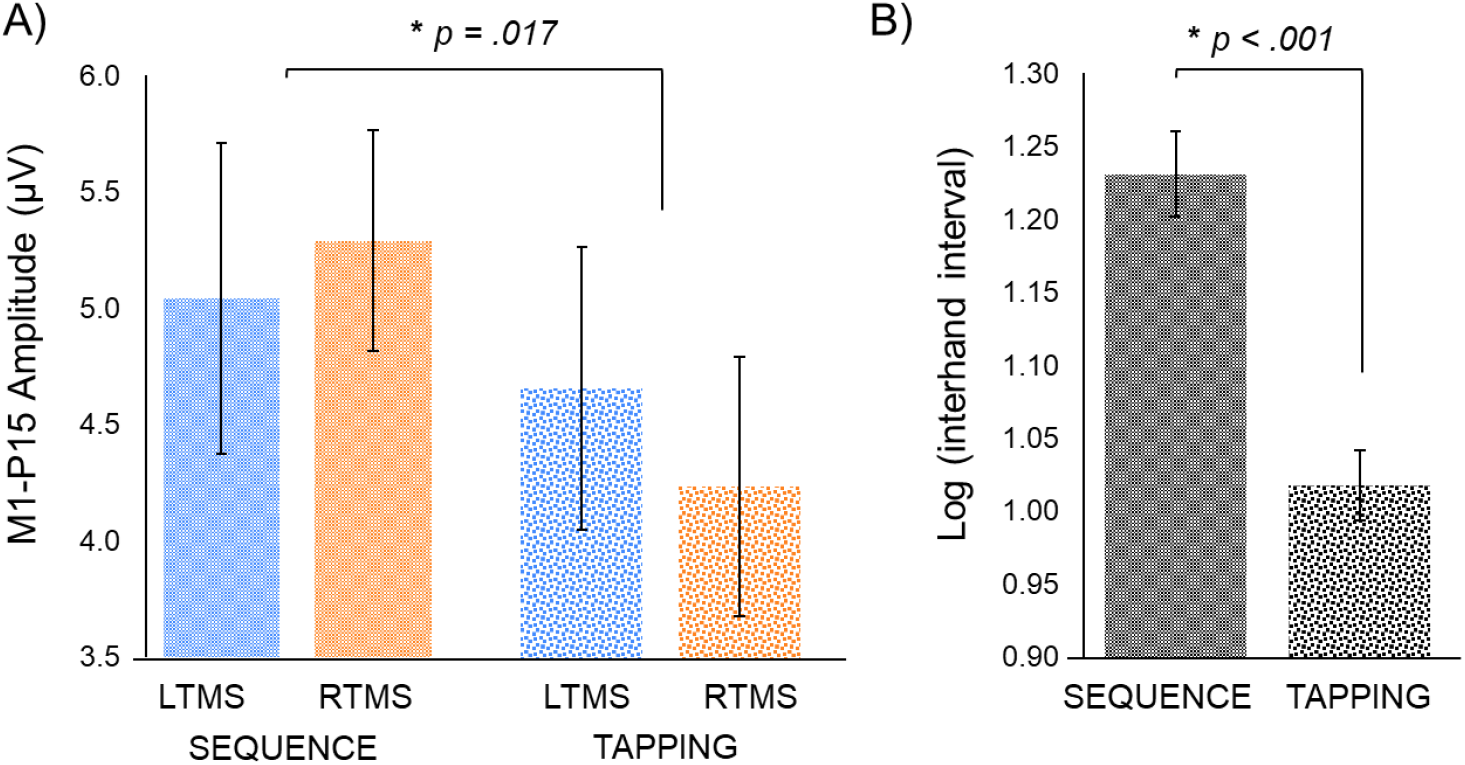
Bimanual task manipulation on M1-P15 amplitude and coordination performance. **A)** Bimanual task affected M1-P15 amplitude, with a larger M1-P15 recorded during the *Sequence* compared to the *Tapping* task (*p* value is referred to the significant main effect of Task in the 2 × 2 rm-ANOVA with factors Hemisphere and Task). **B)** The interhand interval measured during the behavioral *Sequence* task was significantly longer than the one recorded during the behavioral *Tapping* task (*p value* resulting from dependent-samples t-test).

## Discussion

In the present preregistered study we aimed at replicating and extending our previous findings on M1-P15, an early TMS-EEG component recorded on contralateral electrodes about 15 ms after M1 stimulation, which may reflect transcallosal inhibition between motor areas [19]. On a new group of participants and using a different TMS-EEG set-up, we successfully replicated our previous results on the relationship between the amplitude of M1-P15 and the magnitude of the iSP, further corroborating the hypothesis that M1-P15 conveys information on transcallosal inhibition of contralateral M1. In contrast with previous findings, however, the transcallosal conduction delay as indexed by M1-P15 latency was not predictive of bimanual coordination performance. Finally, the present data provides novel evidence of a task-dependent modulation of the strength of transcallosal inhibition as measured by M1-P15 during bimanual movements, but not during an iSP paradigm.

Consistently to our previous findings, M1-P15 could be detected with a comparable topographical pattern across all experimental conditions, after TMS over the left M1 and the right M1 (Figure 2). In our previous work [19], we suggested that the amplitude of M1-P15 represents the inhibition conveyed by the stimulated M1 to its homologous along the fibers of the corpus callosum, by reporting a significant relationship between the magnitude of M1-P15 and of the iSP – a well-known peripheral measure of interhemispheric inhibition [36,37]. In the present study, we provide a replication of this relationship, such that the larger the M1-P15, the greater the normalized iSP area (Figure 2A). Importantly, control analyses ruled out the risk of an artifactual contamination that could have explained the relationship between the two measures, showing that neither M1-P15 amplitude nor iSP normalized area were associated with TMS intensity (Figure 2B-C; [38]). The characterization of early TEP components as reflecting contralateral activation in the motor network is consistent with previous TMS-EEG studies, in which brain source modeling localized the response in the first tens of ms after left M1 stimulation in the right M1 [4,18], and more generally with evidence from double-coil paradigms [36,39]. Nevertheless, the present findings take this body of evidence one step forward, characterizing a specific feature of a TEP component, namely the M1-P15 amplitude, as a reliable cortical measure of the strength of transcallosal inhibition of contralateral motor areas. Indeed, the hypothesis that the iSP is conveyed by transcallosal cortical fibers rather than by uncrossed ipsilateral cortico-spinal pathways has been supported since very early studies, based on the latency iSP [36,40], especially for the APB muscle [41], and on data from patients with-abnormalities of the corpus callosum [20]. Interestingly, it has been recently suggested that the reduced EMG activity observed in the ipsilateral muscles after M1-TMS may not reflect a general and undifferentiated inhibition between motor cortices, but it may arise from a more complex integrative function ensured by mechanisms of surrounding inhibition [42]. Therefore, the same reasoning may apply to M1-P15: although it is associated with an inhibitory effect recorded on ipsilateral muscles, it cannot be excluded that M1-P15 may also subtend narrowed excitatory effects.

Then, we investigated whether we could reproduce the predictive value of M1-P15 latency recorded during the iSP paradigm on bimanual coordination in sequential thumb-to-finger opposition movements. Contrary to our expectations, we did not observe any significant relationship between the M1-P15 latency and behavioral performance at the *Sequence* task (Figure 3). Control analyses on behavioral performance throughout the experimental blocks rule out possible confounding effects of learning on fatigue that may have impacted the performance when considering all blocks together. A possible reason that may explain this null result is that the conditions in which M1-P15 was recorded during the iSP paradigm were not identical to our previous study [19]. Beside the classical iSP condition in which the contralateral hand was at rest [37], in both studies we also recorded a condition in which the contralateral hand was motorically active, expecting to increase the interhemispheric inhibition [43]: while in Bortoletto et al. (2021) participants were involved in visually-cued thumb-to-finger opposition (*Task* condition), in the present study they were asked to maintain a certain level of muscle contraction (*Contracted* condition). The relationship between M1-P15 and bimanual coordination was further investigated by recording TMS-EEG also during the same bimanual movements performed in the behavioral tasks. Results show that M1-P15 was elicited also during the bimanual tasks, with topographical patterns comparable to the ones observed during the iSP paradigm (Figure 2). This finding suggests that the transcallosal information transfer conveys an inhibitory function also during the execution of bimanual movements, supporting the hypothesis that functional inhibition of contralateral M1 may be needed to ensure neural cross-talk at the corticospinal level (but see [42]). Nevertheless, even in this case we did not observe a significant relationship between M1-P15 latency and behavioral performance. It has to be noted that in Bortoletto et al. (2021) the TMS was delivered at the time of movement initiation, which may be a critical timing for M1-M1 interaction, while here it was delivered randomly between one finger movement and the following one. Overall, the discrepancies between this study and our previous one highlight the need for future investigations on the relationship between transcallosal conduction delay and bimanual coordination.

Finally, the aim of modulating M1-P15 according to the state of the motor cortex was fulfilled in an exploratory way. Looking at the bimanual tasks, we found a modulation of the behavioral performance (Figure 5B) and also the M1-P15 amplitude, with a larger M1-P15 amplitude during the sequential (*Sequence*) compared to the repetitive (*Tapping*) bimanual movements (Figure 5A). This result represents the first evidence of a task-dependent modulation of M1-P15, further suggesting the physiological, non-artifactual nature of this early TEP component. Furthermore, the interpretation of this effect suggests that the execution of sequential bimanual movements requires stronger interhemispheric inhibition compared to repetitive bimanual movements. Intriguingly, this hypothesis is in line with evidence on patients with multiple sclerosis [27] and with callosotomy and agenesis of the corpus callosum [44,45], showing that sequential but not repetitive finger opposition movements rely on callosal integrity.

On the other hand, the iSP paradigm was found to be inadequate for the aim, as the iSP was not modulated by hand contraction. In fact, during the iSP paradigm we manipulated the activity of the contralateral hand, expecting to induce a stronger interhemispheric inhibition in the *Contracted* compared to the *Relaxed* condition [43], which we did not observe. The effect of contralateral hand activity on interhemispheric inhibition during the iSP paradigm was absent not only when considering the M1-P15 amplitude and the iSP normalized area combined in the same statistical model, but also when analyzing the two measures separately. The null result on iSP modulation was surprising, considering the consistent effects reported previously throughout several manipulations of contralateral hand activity [43]. However, even in our previous study [19] we did not observe a modulation of iSP depending on contralateral hand activity (no main effect of Task, *p* = 0.86). A possible explanation for these negative findings relies on the quantification method that we adopted to measure the iSP. Here, we employed the normalized iSP area, which has been shown to be more reliable when compared to other measures, such as the iSP area used by Giovannelli and colleagues [43], especially in conditions of different contraction levels [37]. Therefore, further research is needed to understand the nature of this inconsistency. Nevertheless, considering that the magnitude of the iSP and the amplitude of the M1-P15 are expected to reflect the same process, i.e. the strength of interhemispheric inhibition, it is not surprising that even M1-P15 amplitude was unaffected by contralateral hand activity.

## Conclusions

Taken together, our findings support the M1-P15, an early TEP component after M1 stimulation, as a reliable cortical index of interhemispheric inhibition between motor cortices, as revealed by the successful replication of its positive relationship with the iSP. Furthermore, we introduced novel evidence of a task-dependent modulation of M1-P15 amplitude during bimanual tasks, likely depending on the involvement of transcallosal connections for task execution. In future studies, further investigation is required to understand the relationship between transcallosal conduction delay and bimanual coordination, as well as the effects of contralateral hand activity on interhemispheric inhibition during iSP paradigms.

To the best of our knowledge the present study is the first example of preregistration in the TMS-EEG field. While certainly challenging especially for the technical aspects of TMS-EEG coregistration, preregistration of TMS-EEG studies is feasible and thus should be considered in future studies as a powerful strategy to increase the methodological rigor and transparency and to provide an unbiased picture of the results, eventually leading to an improvement of research quality.

## Funding

This article was supported by the Italian Ministry of Health funding ‘Ricerca Corrente’ and by the Department of Philosophy ‘Piero Martinetti’ of the University of Milan with the Project “Departments of Excellence 2018-2022” awarded by the Italian Ministry of Education, University and Research (MIUR) to GB.

## Supplementary Materials

### Pilot experiment

A pilot experiment was performed before data collection with the following aims: P1) ensuring the feasibility of the new finger-sensors to measure behavioral performance in unimanual and bimanual movements, P2) ensuring the presence of M1-P15 during bimanual movements in TMS-EEG recording, and P3) identifying the optimal time range for TMS delivery during bimanual movements. Furthermore, we used the data of two pilot subjects to qualitatively compare the TEPs obtained after high-pass filters at 1 Hz and at 0.1 Hz.

#### Participants

Six young healthy participants (3 women, mean age ± SE: 28 ± 2.1 years, right handed). The same participants were not enrolled in the main experiment.

#### Design and procedure

Participants underwent a single-session within-subject design, including behavioral tasks and a TMS-EEG recording during the bimanual ‘sequence’ task. The behavioral tasks were identical to the one of the main experiment. In the TMS-EEG recording, 20 blocks of single pulse TMS were delivered over the left and the right M1 at 110% of rMT (10 blocks per hemisphere). In each block, TMS was delivered at 30 different time intervals after the metronome sound (from 0 to 480 ms, aligned with refresh rate, steps of 16.7 ms). On average, TMS was delivered every 4 movements; the ISI between TMS pulses was 1 s. Since the randomization of the TMS pulses associated with metronome sounds did not guarantee that the time interval between two consecutive TMS was at least 1 s, we added 2 sounds (i.e., 1 s) between those separated by less than 1s; therefore, the total number of the metronome sounds can slightly vary among blocks and participants. TEPs were analyzed by averaging over 5 time windows (i.e, 0-100 ms, 100-200 ms, 200-300 ms, 300-400 ms, 400-500 ms after the metronome sound), each comprising 6 intervals for TMS delivery, and resulting in a total of 60 TMS pulses for each time window and hemisphere (Figure S1A).

#### Analysis

TMS-EEG data was analyzed following the same pipeline described for the main experiment. On the data of two subjects, we run a control analysis in which the frequency of the high-pass filter was set at 0.1 Hz.

#### Results

Results of the pilot experiment were the following: P1) the recording of the first 3 participants revealed a few technical problems with the new finger-sensors; these problems have been solved for the remaining participants, for which we were to successfully measure a clean signal for both bimanual and unimanual touches with a sampling rate of 9600 Hz; P2) M1-P15, defined as a contralateral positive component peaking ~15 ms after the TMS pulse, was present also during bimanual thumb-to-finger sequential movements (Figure S1B); P3) M1-P15 was present across all intervals of TMS delivery (Figure S1C).

In summary, the pilot experiment enabled us to improve our new finger-sensors and to ensure that M1-P15 is present also during bimanual thumb-to-finger opposition movements independently of the timing of TMS pulses after the metronome sound. Therefore, in the TMS-EEG recording during bimanual tasks of the main experiment TMS pulses were randomly delivered in the time interval between the metronome sounds.

**Figure S1.**
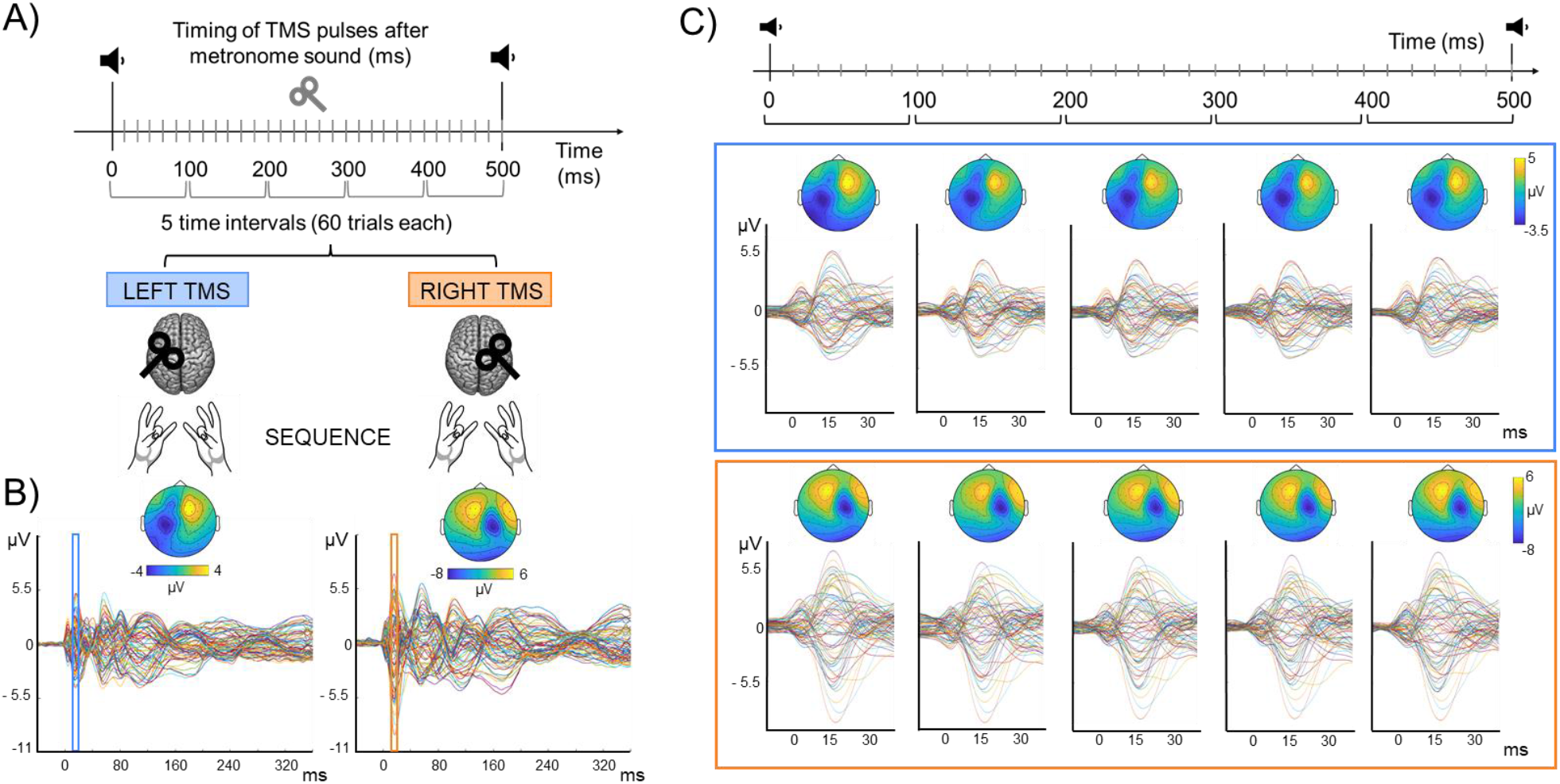
Pilot experiment methods and results. **A)** Schematic representation of the experimental design: while participants were performing the *Sequence* task, TMS was delivered in 30 possible time intervals from the metronome sound, that can be grouped into 5 time intervals of 60 trials each. Both hemispheres were stimulated in different blocks, in counterbalanced order among participants. **B)** Grand average of TEPs and topographical maps after LTMS (left panel) and RTMS (right panel), merging all time intervals. Topographies were obtained by averaging over time between 10 and 20 ms after the TMS pulse; amplitude range is shown in colorbars. **C)** Grand average of TEPs and topographical maps for each of the 5 time intervals of TMS delivery after the metronome sound, for LTMS (upper panel) and RTMS (lower panel), showing no macroscopical modulation of M1-P15 among the different intervals. Amplitude range of topographical maps is shown in colorbars.

**Figure S2.**
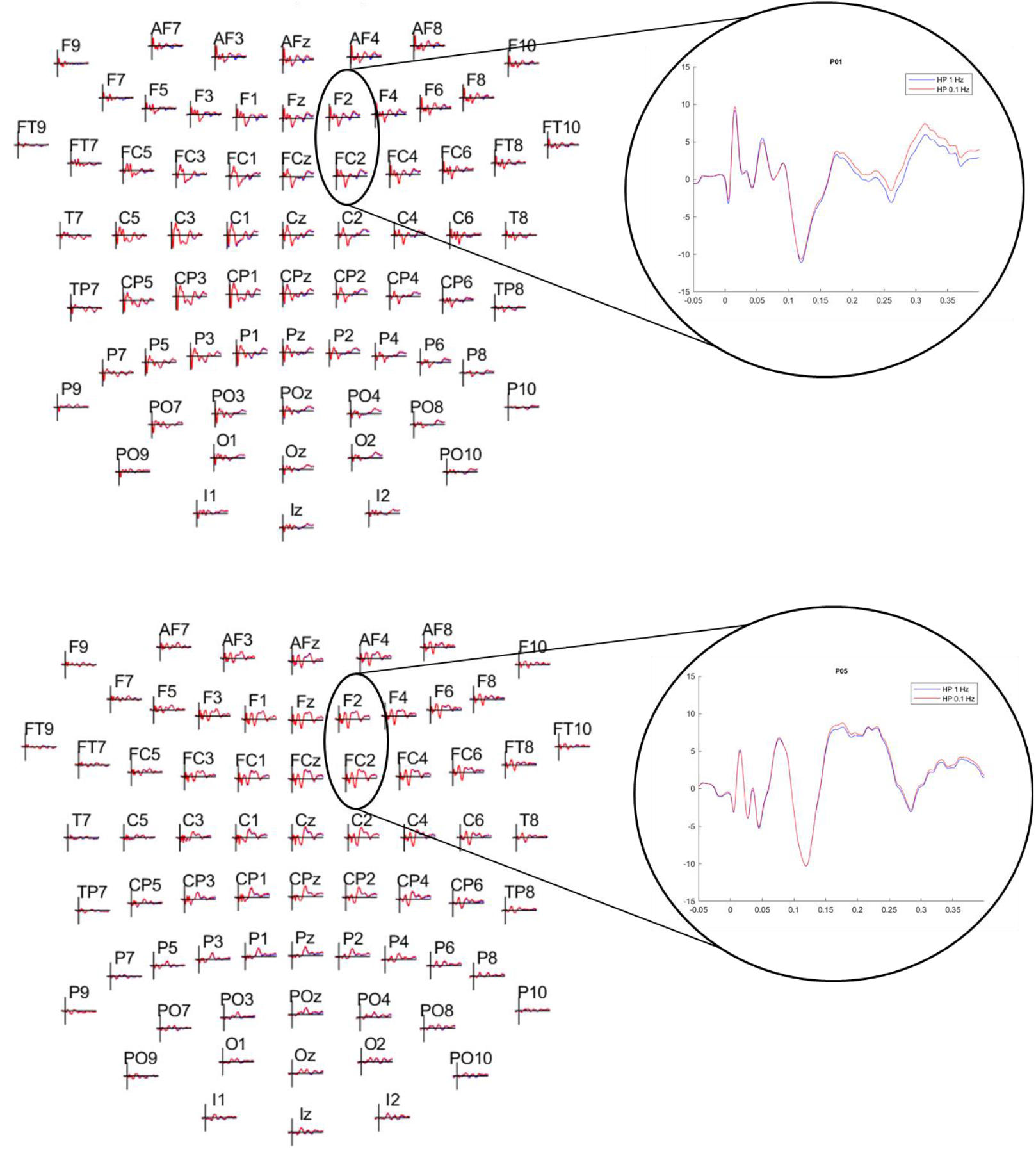
Comparison between high-pass filtering at 1 Hz and 0.1 Hz in two pilot subjects. Average over trials showing the overlay of TEPs obtained after 1 Hz filter (blue trace) and 0.1 Hz filter (red trace). The enlargement on the right shows the average signal between the two electrodes considered for M1-P15 after LTMS (i.e., F2, FC2).

### Main experiment

#### Unimanual task

In the unimanual motor task, participants were asked to tap as fast as possible with their index finger on a sensor placed on the desk within a time interval of 10 s, while fixating a cross in the center of the screen. The ‘go’ signal was represented by the onset of a white noise, delivered through earphones. In different blocks, they were asked to use their right and left index finger, respectively, performing 2 runs for each hand; hands order was counterbalanced across participants.

## Notes

### Competing Interest Statement

The authors have declared no competing interest.

